# Habit learning shapes activity dynamics in the central nucleus of the amygdala

**DOI:** 10.1101/2024.02.20.580730

**Authors:** Kenneth A. Amaya, J. Eric Carmichael, Erica S. Townsend, Jensen A. Palmer, Jeffrey J. Stott, Kyle S. Smith

**Author notes:** Corresponding authors: Kenneth A. Amaya, Kyle S. Smith.

## Abstract

As animals perform instrumental tasks, they may develop a habit response with extended experience. Habits are automatic, inflexible, outcome value insensitive behaviors that are regulated by a network of brain regions including the central nucleus of the amygdala (CeA). Prior work has demonstrated that the CeA governs motivational pursuit and is necessary for habit formation. However, the behavioral features that CeA neurons encode in habit formation remain relatively unknown. To address this, we first used male and female Long-Evans rats to quantify CeA cFos expression after performance of a maze task. There, we found that animals with extended training show elevated cFos expression. Then, we implanted animals with drivable silicon probes to record *in-vivo* single unit electrophysiological activity from the CeA as animals developed habit responding on the maze. We observed significant activity during outcome consumption late in training while also observing elevated unit activity when animals consumed outcomes of larger magnitudes. Outcome related activity did not persist during probe tests following outcome devaluation, despite animals continuing to perform the task. Together, these data add to growing evidence that suggests that the CeA is involved in motivational processes that contribute to the development of habit formation.

## Introduction

When repeatedly performing instrumental tasks, animals can transition from a reliance on goal-directed behaviors that are contingent upon associations between actions and outcomes, to habits, governed by associations between responses and antecedent stimuli ^1^. Distinct cortical, striatal, and midbrain (dopaminergic) contributions have been identified that contribute to these two learning processes ^2–7^. In addition, there is evidence that the amygdala plays a key role in dictating the balance of goal-directed versus habitual strategies of behavior. Lesions to the central nucleus of the amygdala (CeA) abolish habit formation, as does functionally disconnecting the CeA from the dorsolateral striatum (DLS) ^8^. Stimulation of the CeA can increase motivated behaviors to the point of compulsion ^9–11^. Recent work has noted that the CeA and its outputs are preferentially engaged during stress-induced accelerated habit formation and in potentially addictive behaviors like ethanol-seeking ^12,13^. However, unlike research on other habit-promoting brain areas ^14,15^, little is known about what changes in CeA electrophysiological activity relate to habit learning under normal conditions.

Among studies conducting single-unit electrophysiological recordings in other nodes of the habit network (infralimbic cortex IL, DLS, and substantia nigra SNc), one common population-level activity motif related to habits is action “chunking”. Chunking dynamics are described as the learning-related organization of neural activity such that spiking is at its greatest at the initiation and/or termination of an acquired action sequence ^14 – 18^. Action chunking in several brain areas aligns with the formation of habits defined by vigorous and outcome-insensitive performance ^19^. The development of chunking activity as it pertains to the animal’s behavior varies by brain structure, with DLS chunking emerging earlier in training than it does in IL ^14^. Further, these patterns correlate with different behavioral phenotypes thought to contribute to the overall psychological-behavioral repertoire indicative of a habit, suggesting that while this motif arises throughout the hypothesized habit network, each node of said network may be contributing a unique component of what makes a behavior a habit ^14^. Beyond chunking activity patterns, habit learning has been linked to changes in outcome representations in the DLS ^19^, and also to changes in the DMS activity and its innervation from midbrain and cortical sources ^20–23^. With these studies having established ways in which neural activity changes may encourage habit formation, we sought to examine how changes in CeA activity might do so as well.

## Results

We first trained Fos-GFP+ Long Evans rats (N = 6 male, 6 female) on a custom plus-maze task where they were required to turn a certain direction (right or left) to earn food outcomes available at the ends of the maze arms (Fig 1A). Outcomes varied in identity and magnitude across the four maze end-arms. Animals were allowed 60 minutes per session to complete 100 trials, regardless of whether the trials were correct responses. Experimental groups included animals that were trained to criterion (Group Criteria; 60% correct response criteria for 3 days), were over-trained (Group Overtrained; 7 additional days), or a No-Run control group that was handled similar to the animals in Group Overtrained. Animals in both groups readily and similarly learned this task, as shown by a significant effect of Session (OR: 1.37; CI: 1.21 – 1.54; p < 0.001), but no effects of Group (OR: 1.13; 95% conf. interval (CI): 0.59 – 2.17; p = 0.71) nor was there a significant interaction between the two variables (OR: 0.90; CI: 0.79 – 1.01; p = 0.07) (Fig 1B). Shortly after completion of their final training session (C3 for Group Criteria; O7 for Group Overtrained), animals were sacrificed, and tissue was processed to visualize cFos expression in the CeA (Fig 1C). As shown by a Wilcoxon rank-sum test, no difference in cFos expression was observed between the No-Run control and Criteria groups (W = 12, p = 0.312). However, the Overtrained Group showed significantly elevated cFos expression compared to the control animals (W = 0, p = 0.008) (Fig 1D). No differences in DAPI counts were observed between groups, indicating that the greater cFos expression observed was not a product of more neurons in the field of view (W = 8, p = 0.203) (Fig 1E). These data showed that extended training on the maze task engaged the CeA.

**Figure 1.**
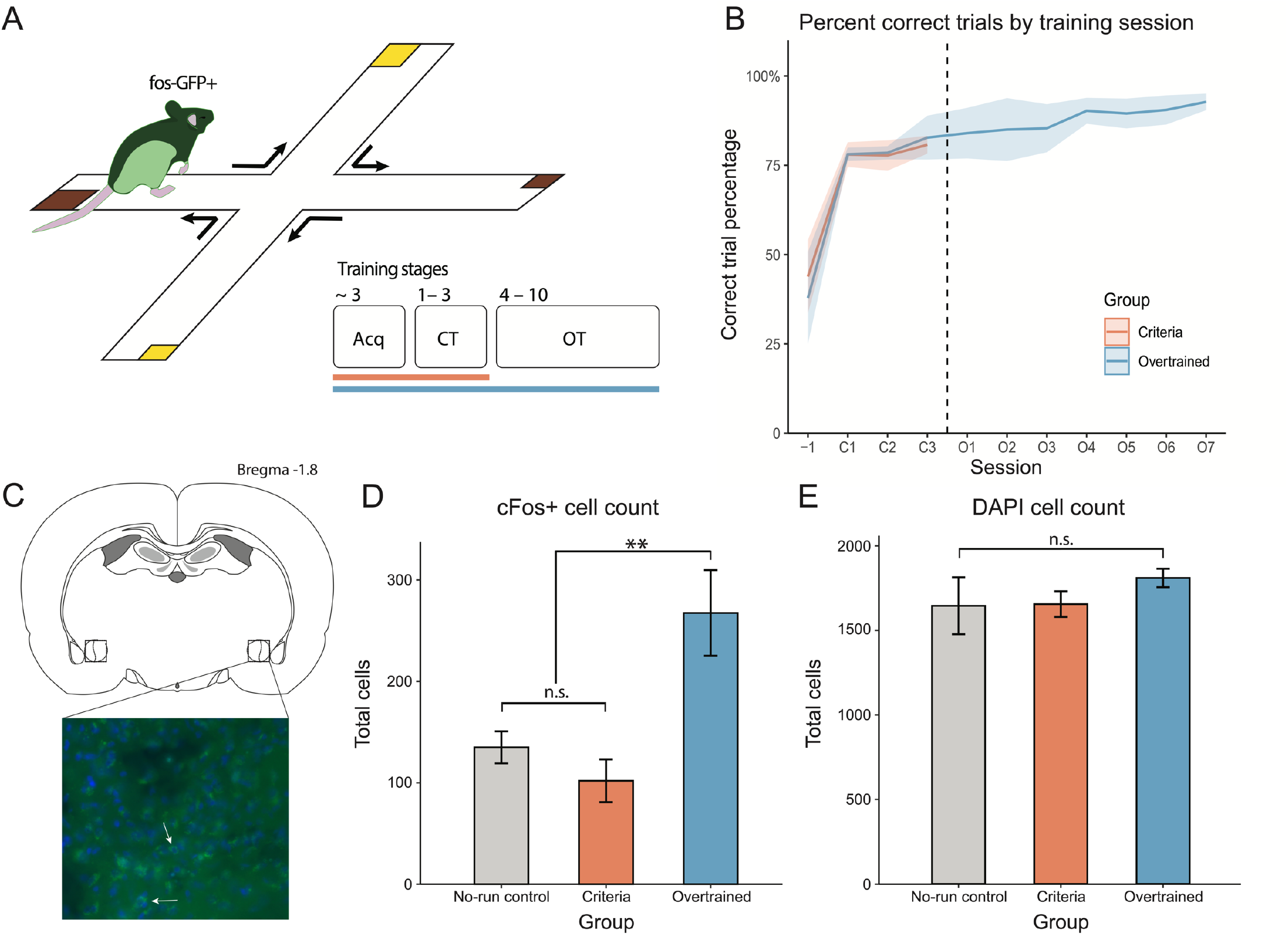
Overtraining increases cFos expression in the central nucleus of the amygdala. A) Animals were trained on a plus maze task where performance of a turn rule (e.g. turn right) resulted in delivery of a food reward (two flavors: banana or chocolate flavored sucrose pellet, two magnitudes: 1 or 3 pellets). B) Animals (N = 8) were divided into two groups, a Criteria-trained group and an Overtrained group that received 7 training sessions more than their Criteria counterparts. C) Animals were sacrificed 90 minutes after the conclusion of their final training session (C3 for Criteria or O7 for Overtrained rats) to allow for the expression and subsequent quantification of cFos. D) Expression of cFos was significantly elevated in the Overtrained group only as compared to the Criteria group and a No-run control group that received similar handling but never experienced the maze. E) DAPI expression amongst the groups was equivalent, indicating that cFos expression changes were not a product of field of view differences.

Next, we implanted silicon recording probes targeting the CeA in Long-Evans rats (N = 5 male, 4 female) and recorded extracellular action potentials throughout training and over-training on the maze task (Fig 2A, 2B, Methods). The task was identical to the prior experiment except outcomes were two distinctly flavored sucrose solutions to reduce eating/chewing noise artifacts in the recordings. Outcomes again varied in identity and size across the maze’s four end-arms and included large magnitude (0.3 mL) grape solution, large magnitude orange solution, small magnitude (0.1 mL) grape solution, or small orange solution. Again, animals readily acquired the task, reflected by their performance above the experimenter-set 60% correct criteria for the duration of the experiment, and a significant effect of session (OR: 1.12; CI: 1.10 – 1.14; p < 0.001) (Fig 2C), a significant effect of session on correct trial rate (est: 0.38; CI: 0.31 – 0.46; p < 0.001) (Fig 2D), and the significantly increasing their speed each session (est: 0.55; CI: 0.23 – 0.86; p = 0.001) (Fig 2E). After completion of the final overtraining session (O7), animals underwent outcome devaluation, in which one reward flavor (e.g., orange) was paired with lithium chloride-induced nausea until that outcome was rejected by the animals (Methods). In a subsequent post-devaluation session under extinction conditions, animals continued making correct responses, suggesting they were habitual (Fig 2F). A series of post-devaluation reacquisition sessions in which rewards were made available again followed. Animals were required to make the same turn responses as before and were thus required to approach the devalued goal in order to trigger a run option to the valued goal; in these conditions, animals maintained task performance and rejected the devalued (but not non-devalued) rewards (Fig 2F-G). There was no effect of Session on correct responding after devaluation (OR: 1.04; CI: 0.92 – 1.17; p = 0.536).

**Figure 2.**
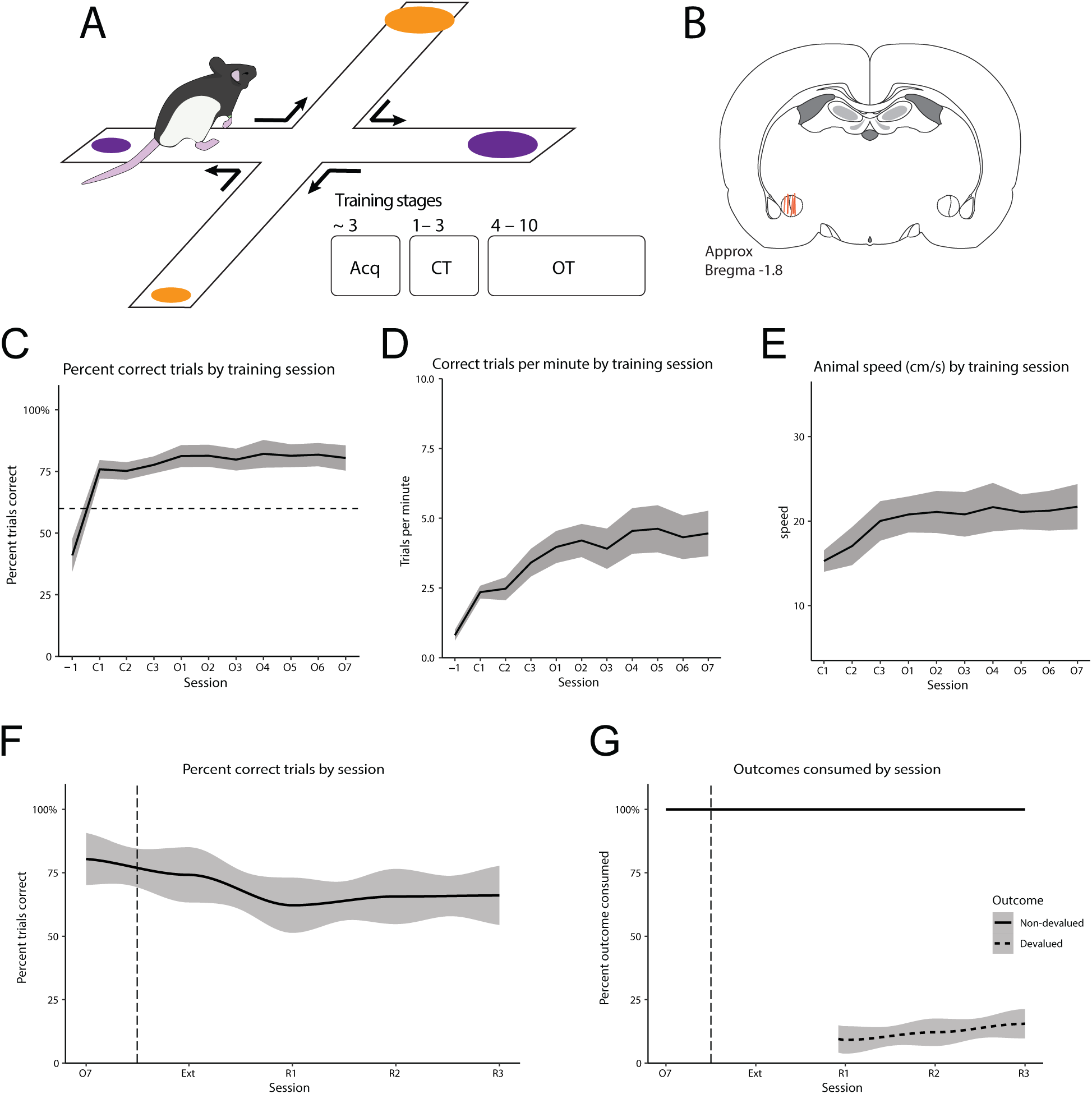
Behavioral results from the recording experiment. A) Animals were trained on a plus maze task similar to that described in Figure 1. Animals (n = 9) were required to turn a certain direction to earn outcome delivery, with unique outcome magnitudes (0.3 or 0.1 mL) and identities (orange or grape flavored 20% sucrose) delivered at each arm end. Training consisted of brief acquisition training, pre-implant, followed by Criteria training and Over-training before animals advanced through outcome devaluation and subsequent post-devaluation behavior probes. B) Silicon probes were implanted targeting the central nucleus of the amygdala. C) Percentage of correct trials performed by session, plotted as the mean with shading representing the standard error of the mean (SEM). The dashed line reflects a 60%-correct learning threshold. D) Correct trial rate by training session, plotted as the mean +/- SEM. E) Mean speed over training sessions with SEM. F) Post-devaluation task performance, plotted as mean percentage of correct trials by session. Sessions to the right of the dashed line are post-devaluation extinction (E1) and reacquisition (R1 – R3, rewarded). G) Mean percentage of outcome consumed by post-devaluation session (no rewards available in Extinction).

Recorded neural activity in the CeA was dynamic during training with populations of neurons showing modulation by action performance and outcome consumption. We recorded 187 isolated units in the CeA with an average firing rate of 6.439 Hz and used spike timing (burst index and firing rate) and waveform properties (spike width) to classify the recorded units into two putative types: pyramidal-like (n = 181) and interneuron-like (n = 6) (Fig 3B). We report a diverse activity from the recorded units (Fig 3D), including both increases and decreases in unit activity during distinct periods of trial completion on the maze task, such as during outcome receipt/consumption (Fig 3A top, bottom) or action performance (Fig 3A middle). To visualize unit responses over time, we created a peri-event time histogram of z-scored firing rates for each of the recorded units across sessions, where time = 0 is the entry of the animal into the reward zone, denoted by the black vertical line (Fig 3C). Red dashed lines indicate the time windows used to analyze unit activity and were approximately tied to behavioral events including trial initiation (t = -3.0), turn execution (t = -1.0), reward zone arrival (t = 0), and the completion of reward consumption (t = +2.0). Our cFos analyses, as well as prior findings^14,17^, suggested that the transition from training through overtraining is when behavior transitioned into habitual control and engaged changes in CeA activity. Therefore, we grouped units by learning phases for subsequent analyses: Criteria (C1 – C3), Early Over-training (O1 – O3), Late Over-training (O4 – O7), and Post-devaluation (R1 – R3).

**Figure 3.**
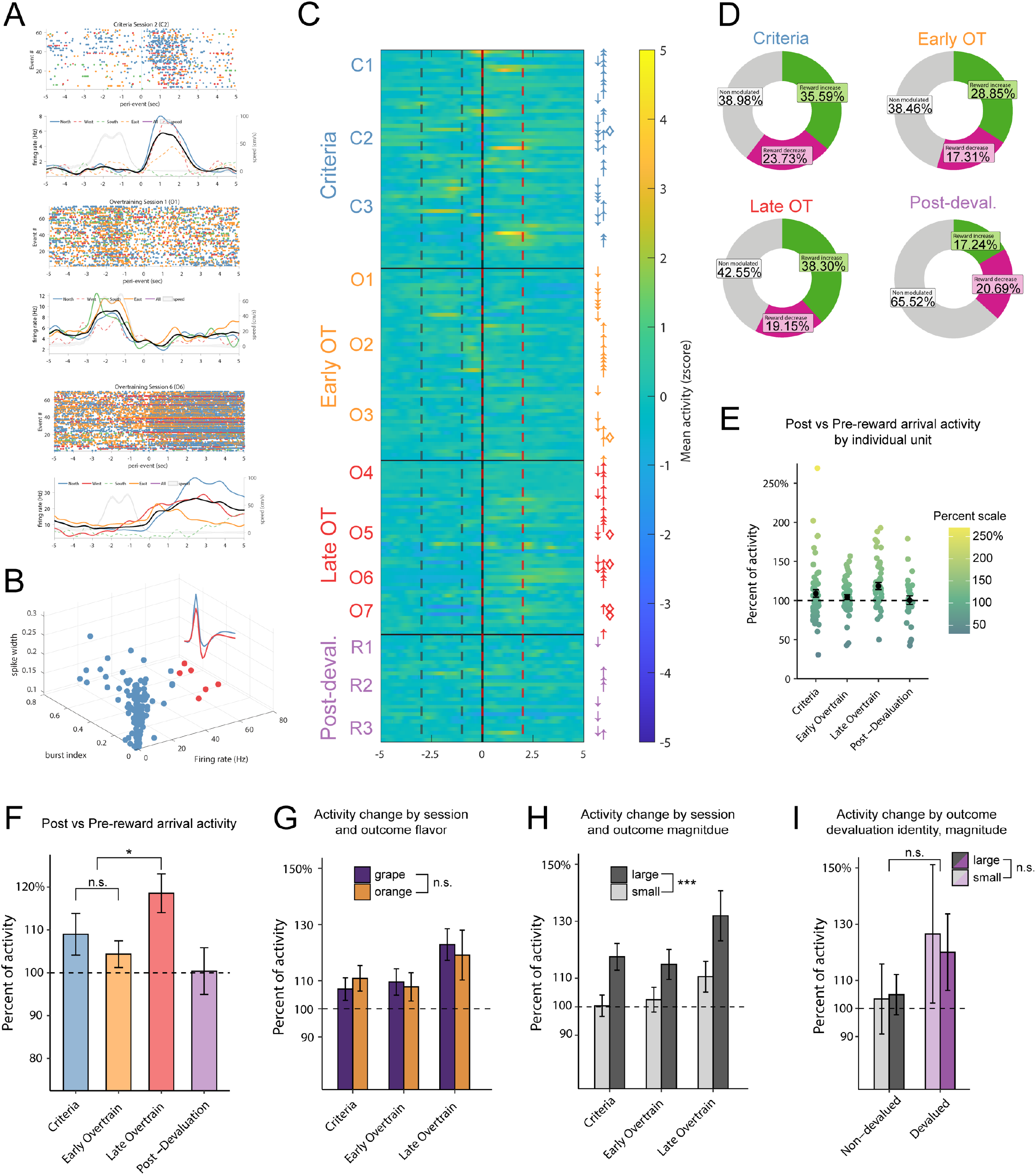
Characterizing behavioral correlates of central amygdala activity during habit formation. A) Example peri-event time histograms from units at different stages of training. Top to bottom: C2 (outcome), O1 (action), O6 (outcome). Colors represent responding on each maze arm (different outcome identities and magnitudes), overall responding shown in Black. Significantly modulation of unit activity is denoted as a solid line, non-modulated activity shown with a dashed line. B) Cell classification of recorded units into two major cell types: pyramidal-like (n = 181, left/blue) and interneuron-like (n = 6, right/red). C) Normalized (baseline-subtracted z-scores) activity of CeA units, constructed from abutted +/- 200 ms peri-event periods during Criteria training, Over-training, and Post-devaluation probe tests. Arrows indicate whether units increased or decreased their responding, diamonds represent putative fast-spiking interneurons. Color scale shown on the right. D) Proportions of units that were non-modulated (gray), whose activity increased in the reward zone (lime green), or whose activity was decreased in the reward zone (magenta), separated by training phase. E) Scatterplot of neural activity taking place around reward zone arrival by training phase. Black dots represent the mean percentage of activity with +/- SEM error bars. Units with values above 100% are reward biased, units with values below 100% show greater activity prior to reward arrival. F) Mean +/- SEM unit activity percentage scores by phase. Data from Fig 3E presented as bar graphs. No difference between Criteria and Early Overtraining responding, significant outcome bias observed in Late Overtraining responding (* p = 0.018). No outcome bias observed in during Post-Devaluation. G) Recorded CeA units do not show biased responding to flavor identity (purple: grape; orange: orange). H) Recorded CeA units are more responsive to outcomes of greater magnitude during all training phases (*** p < 0.001). I) Responding to non-devalued and devalued outcomes grouped by outcome magnitude during Post-Devaluation reacquisition sessions, no difference observed between devaluation identity (p = 0.336), or magnitude (p = 0.443).

Within the myriad of different CeA units responding to different aspects of task performance, the response type that changed to match habit formation from criterion training to overtraining, and then habit decay during post-devaluation sessions, related to outcome signaling. Some units exhibit elevated activity prior to the animal entering the reward area while others show elevated activity while animals were in the reward area, which was calculated as a ratio of firing rates before and after reward area entry (Fig 3E). With this approach, we observed that in Late Over-training the CeA units become collectively more reward-tuned than during training phases, as revealed by a Wilcoxon rank-sum test (W = 1986.5, p = 0.018). This data is shown as a mean of percent activity change with all units displayed by training phase (Fig 3E) and again with recorded units pooled (Fig 3F). We further analyzed outcome-responsive units with respect to outcome identity and size. Different outcome identities (i.e., orange vs. grape) did not modulate the activity of this CeA unit population (W = 50226, p = 0.205) (Fig 3G). However, CeA units did respond more to outcomes of greater magnitude (Fig 3F; W = 39190, p < 0.001) (Fig 3H). This putative sensitivity of CeA units to outcome value was observed at all training stages except the post-devaluation sessions (Fig 3I). During post-devaluation sessions, no responsiveness was observed to reflect outcome identity (devalued vs non-devalued; W = 1203.5, p = 0.336) nor magnitude (W = 1233.5; p = 0.443). However, compared to pre-devaluation sessions, the percent of units that exhibited increased activity to outcome delivery dropped by roughly half (Fig 3D), to the point of being too low a sample size to draw major conclusions from regarding reward value representation.

## Discussion

Here, we report that activity in the central nucleus of the amygdala (CeA) changes dynamically with habit formation. Greater cFos expression in the CeA was observed after extended training on the plus-maze, aligning with prior studies that detail a critical role for the CeA in habit formation ^8,12^. Through *in-vivo* single-unit recordings, we show that CeA neurons are modulated by specific task-related events on a plus-maze task. The pooled activity of neurons shift responding as training progresses to accentuate the reward outcome events of the task. While no explicit chunking motifs were observed, the emphasis on the outcome event in CeA activity emerged late in training, as habits formed, and then decayed after reward devaluation. These results add to a growing body of literature implicating the CeA in appetitive learning processes, including habit formation ^8,12,24–26^, and add confirmation to recent work showing reward-related changes in CeA activity that reflect reward preference ^13^ as well as reward-related signals that reflect stress-facilitated habit-learning in the CeA-to-dorsomedial striatum (DMS) pathway ^12^.

Increased outcome representation in the CeA as a habit develops might be unexpected from concepts of habit learning. Habits are thought to be guided by associations between stimuli and the responses they elicit. Therefore, in the brain, signals for habits would occur at the times of action selection and execution. Habit-related activity occurring during actions has been observed in many brain areas, notably in the dorsolateral striatum where neural activity grows to emphasize action initiation and the strength of this activity can dictate how habitual behaviors are ^14,17–19,23^. As loss of CeA function produces seemingly similar consequences on behavior as loss of dorsolateral striatum (DLS) function in this domain ^8^, namely less habitual behavior using measures like sensitivity to reward devaluation and action-outcome degradation, it is notable that the CeA representation of habits is unlike that seen in similarly habit-promoting areas like the DLS. However, aside from DLS representation of actions, there are a small number of DLS neurons that respond to rewards (or lack of rewards) ^27^. These DLS responses to outcomes also change over the course of habit learning, such that they grow to become less reward specific, and such that their response to lack of reward declines ^19^. Thus, there may be some relation between DLS and CeA in reward-related processes that support, in some manner, habitual behavior. Although increased reward representation may be surprising given that there is no outcome representation involved in guiding habits (at the time of action performance) ^3,28^, the outcome occurring after a habit may act as a reinforcer, strengthening the relationship between the stimulus and response ^3^. Presently, it is possible that the observed bias in neuronal responding in the CeA that emerges as animals train on the maze task represents strengthening of the running pattern that was just performed.

Although stimulus-response-(reinforcement) learning may well reflect how the CeA contributes to persistent behaviors such as the devaluation-insensitive maze running here, there is also the possibility that it does so through a different process of reward saliency. In several studies, it was found that optogenetic stimulation of CeA neurons at the time of reward delivery leads to highly perseverative reward seeking, to the extent that it resists large levels of aversive shock paired with the reward ^10,11,29^. Further, pairing CeA stimulation with an aversive outcome can even endow it with motivationally attractive qualities ^9^. This striking persistence in motivation to receive the CeA-stimulation-paired outcomes could reflect an underlying function for the CeA in tagging reward events with motivational salience. In this view, our finding that reward-related activity grows in the CeA as behaviors become over-trained and habitual, which is also observed when stress exposure exacerbates habit formation ^12^, could cause behavioral persistence because of increased motivational value of that reward event. By extension, it is possible to consider that some forms of habit-like behaviors, defined by insensitivity to aversive shock or insensitivity to reward devaluation, could result from a motivational saliency process rather than a stimulus-response-(reinforcement) learning process.

Further concerning motivational functions of the amygdala, the basolateral amygdala, which sits adjacent to the CeA, has been implicated in the use of outcome-specific representations to guide action during Pavlovian-to-Instrumental transfer (PIT) tests. In other words, the BLA is required for specific-PIT effects to be observed, where animals will preferentially perform Action1 to earn Outcome1 upon hearing Cue1, despite never experiencing that Cue in conjunction with access to the manipulandum to perform the action ^26^. The inferential nature of PIT is intriguing to use as a model for motivation, especially as a means to gauge how animals are using outcome representations to guide their behavior. While the BLA is necessary for specific-PIT, the CeA is not, as lesioning the CeA has no impact on an animals ability to perform the task ^26,30^. However, deficits were observed in general PIT, where outcome representations were required to guide behavior, suggesting that the CeA is more involved in general appetitive motivation rather than outcome identity encoding ^26^. Presently, the recorded CeA activity appears to support these notions regarding what the nucleus may be associated with behaviorally. We do not observe any bias in responding to one flavor over another, suggesting that this information is not necessarily represented. Importantly, we do observe a robust bias in responding to outcomes of greater magnitude. It is possible that the CeA encodes value or salience, putatively part of the value computation, which the animal can then use to promote appetitive seeking behaviors.

Understanding the networks that govern habit formation and expression is dependent on integration of the presently observed CeA activity patterns with activity in other habit nodes. While further work will be needed to parse the contributions of specific CeA populations to habit formation, our present characterization offers an opportunity for speculation. Primarily, recent work has shown that projection neurons from the CeA to the substantia nigra can promote appetitive associative learning ^24^. Given that the substantia nigra pars compacta (SNc) sends habit-relevant dopaminergic projections to the DLS ^31^, a semblance of a network appears to be forming, with CeA inputs regulating SNc activity that subsequently contacts the DLS. Importantly, the CeA is primarily composed of GABAergic projection neurons ^32^, which presents the possibility that enhanced CeA activity promotes habit formation and expression through inhibition of local regulatory activity in the SNc. Excitingly, a recent report states that the CeA projects preferentially to local GABAergic neurons in the substantia nigra ^24^. Thus, this circuit could be potentially involved presently during habit formation on our task. An alternative to this hypothesized network comes from a report that highlights CeA projections to the DMS that govern habit formation after animals undergo a chronic unpredictable stress protocol ^12^. There, experimenters used fiber photometry to show elevated CeA calcium activity following outcome collection, but only in animals that had undergone a stress protocol and were thus reliant on habit responding. Our current results focus on the CeA single-unit electrophysiological activity generally, not on any specific projection population, leaving the door open on whether our recorded units were part of either of these pathways. Future work should pay careful attention to the networks that the CeA is activating to facilitate habit formation in both healthy and disordered states.

## Methods

### Subjects

Individually housed transgenic cFos-GFP Long-Evans Rats (N = 12) and Long-Evans Rats (N = 9) were maintained on a reverse light-dark cycle and within 85% of their pre-surgical weight. The cFos-GFP transgenic rats were provided by the laboratory of Dr. Bruce Hope (NIDA). Animals were run in experiments during their dark (active) cycle. All procedures were in accordance with the National Institute of Health’s Guide for the Care and Use of Laboratory Animals. Protocols were approved by the Dartmouth College Institutional Animal Care and Use Committee.

### cFos experiment behavior

Training began approximately 1 week following the commencement of food deprivation, once animals were at the target weight threshold of 85% of their *ad libitum* weights. Animals were split into three groups (n = 4 per group): Group Criterion, Group Overtrained, and Group No-Run. The No-Run group experienced the same handling techniques but no behavioral procedures to control for the frequent handling and backpack-wearing of experimental groups.

Groups Criterion and Overtrained first underwent an initial acclimation session on a custom plus-shaped maze in a dimly lit room with ambient white noise playing (∼20 dB) while wearing LED backpacks where chocolate-flavored and banana-flavored grain pellets were freely available in the food sites (Bioserv #F0299 and #F0059, Flemington, NJ). The next day, animals began maze training where a specific response (turn right or turn left) was required to earn food outcomes at each arm end. The response requirement remained the same for each animal throughout testing but was counterbalanced across animals. Accurate entry into end-arms of the maze resulted in reward delivery that was triggered by entry into the end-arm. Rewards were different for each end-arm. One arm contained large magnitude (3 pellets) of chocolate, another large magnitude of banana, another small magnitude (1 pellet) of chocolate, and the last small magnitude banana. Training for Group Criteria occurred until three consecutive sessions at 60% correct responding were met. Training for Group Overtrained extended for an additional seven sessions beyond what Group Criteria completed, totaling 10 completed sessions at or above 60% accuracy.

### cFos experiment histological procedures

Animals were sacrificed 90 minutes following the completion of the final training session for each group to allow for expression of the immediate early gene cFos. To accomplish this, animals were deeply anesthetized via isoflurane then transcardially perfused with 0.9% saline followed by 10% formalin. Brains were extracted and stored in 20% sucrose for 2 days before being sectioned at a thickness of 60 microns, targeting the central nucleus of the amygdala at the coronal plane 1.8mm behind bregma, informed by prior literature ^8^. Sections were mounted onto microscope slides and cover-slipped with DAPI-containing hard set mounting medium (Vectashield; Vector Laboratories, Burlingame, CA, USA). Bilateral GFP expression and DAPI labeling in the CeA was visualized via fluorescence microscopy and images were saved for cell counting.

### cFos experiment data analysis

Animal learning was analyzed using generalized linear mixed models to model Correct Completion Percentage as predicted by Group, Session, and the interaction between Group and Session. Odds ratios, 95% confidence intervals, and p-values are reported. Counts of cells expressing cFos were collected in ImageJ by an experimenter blinded to the group identities of the animals. DAPI-labeled cells were also counted by the same experimenter and overlap between the two counts were used to verify that putative cFos-labeled cells were indeed neurons. Data analysis of cell counts proceeded by comparing expression of cFos+ cells in the CeA between groups using a non-parametric test, the Wilcoxon rank sum test with continuity correction, as count data is not normally distributed. These methods were replicated for the DAPI count data. Analyses and figure generation were conducted in R (R Core Team 2016, “stats”, “lme4”, “lmerTest”, “car”, “ggplot2”). Figures were stylized in Adobe Illustrator.

### Recording experiment surgery

Rats used in the electrophysiological recording experiment were implanted with 16-channel, 50-μm-thick silicon probes (A2×2-tet-10mm-150-150-121; NeuroNexus, Ann Arbor, MI). Probes were ordered attached to a chronic microdrive (dDrive XL, NeuroNexus) that stood 15.5 mm in height and featured a maximum driving distance of 7.0 mm. During surgery, rats were anesthetized via isoflurane (5% induction, 1 –3% maintenance) and a non-steroidal anti-inflammatory agent was administered pre- and post-operatively to reduce pain and discomfort. Silicon probes were implanted unilaterally over the central nucleus of the amygdala (relative to bregma AP: -1.8 mm; ML: +4.0 mm, per Lingawi and Balleine 2012) and lowered via the stereotaxic arm until the base of the drive mechanism made contact with the surface of the skull (typical probe depth about 2.4 mm into the brain). Once at this resting depth, the drive mechanism was affixed to the surface of the skull via Metabond dental cement (Parkell, Edgewood, NY) and implanted bone screws that held down copper mesh, to be used later in the surgery to create an on-head faraday cage to reduce noise and protect the implant. A ground screw (over cerebellum, contralateral to implant site) and a reference screw (over cerebellum, ipsilateral to implant) were then connected to the ground and reference wires coming from the probe and cemented into place. Probes were lowered approximately 3.0mm further during the surgery (final surgical probe depth: 5.2 – 5.6 mm). The on-head faraday cage was constructed and reinforced with dental cement and mated with a custom-made cap to allow for future access to the probe. Surgical procedures were adopted from past literature ^33^. Following surgery, animals were given post-operative intraperitoneal injections of 0.9% saline and the antibiotic Baytril for three days. Probes were lowered to the target depth (7.8 mm deep) over the course of 10 post-surgical days and were subsequently moved in < 0.02 mm steps.

### Recording experiment behavior

Separate animals were implanted with electrodes into the CeA. Maze training began before surgical implantation, approximately 1 week following the commencement of food deprivation, once animals hit their threshold of 85% of their *ad libitum* weights. Animals first underwent an acclimation session while wearing LED backpacks where grape-flavored and orange-flavored 20% sucrose solution was freely available at the food sites. Outcomes were flavored 20% sucrose solutions to eliminate chewing noise artifacts noise detected by the probes. The same magnitude differences were used at the ends of distinct arms in both experiments. Importantly, outcome type (pellet-cFos or solution-ephys) was consistent within experiments.

Animals then began maze training where a specific response (turn right or turn left) was required to earn outcomes at each arm-end. Task rules (right versus left turn) were counterbalanced. The four arm ends delivered either large magnitude (0.3 mL) grape solution, large magnitude orange solution, small magnitude (0.1 mL) grape solution, or small orange solution for correct responses. Pre-surgery training was completed once animals finished 100 trials (including either correct and incorrect response types) within the allotted 60-minute session time limit. Surgical implantation of silicon probes followed pre-training.

Training resumed approximately 1.5 weeks following surgery, once animals hit their threshold of 85% of their *ad libitum* weights. Animals resumed maze training. Training proceeded through overtraining (3 sessions at Criteria plus 7 sessions of overtraining) for all animals. Following overtraining, animals underwent outcome devaluation by conditioned taste aversion, where free access to one reward flavor was given on two of the maze arms (North/South or East/West) and paired with a 0.3M injection of lithium chloride (10 mg/kg) to devalue one of the two reward identities (grape or orange). Devaluation proceeded until animals rejected the rewards delivered, as previously described ^34^. Following devaluation, a brief 5-minute extinction probe session was conducted in which animals were free to perform the maze task but no rewards were delivered. Subsequent reacquisition probe sessions were then administered where animals could solve the maze (same rule as before devaluation) and correct responses were again rewarded. Post-devaluation reward consumption was recorded during the reacquisition sessions.

### Recording experiment behavioral data analysis

Animal learning was analyzed using generalized linear mixed models to model Correct Completion Percentage as predicted by Session. Odds ratios, 95% confidence intervals, and p-values are reported. For measures of completion rate and animal speed, linear mixed models were used to model those variables as predicted by Session. There, effect estimates, 95% confidence intervals, and p-values are reported. Analyses and figure generation were conducted in R (R Core Team 2016, “stats”, “lme4”, “lmerTest”, “car”, “ggplot2”). Figures were stylized in Adobe Illustrator.

### Neuronal spiking data acquisition and processing

Electrical signals from silicon probes were recorded via a head mounted HS-18 preamplifier and synchronized with behavioral tracking and task events using a Digital Lynx SX acquisition system (Neuralynx). The data was sampled at 30kHz, filtered between 600-6000Hz, and spike detection thresholds were set on a tetrode-by-tetrode basis before the animal began the task. When a spike was detected, 32 samples of the continuous signal representing the spike waveform were saved for offline sorting in MClust 4.3 (A. D. Redish et al.). Spike isolation was initially automated using KlustaKwik (K. Harris) before manual refinement and curation. Only neurons with sufficient firing over the entire recording session (>0.2Hz) were included in the analyses. Neurons were further classified into putative fast-spiking and regular-firing types using k-means clustering based on spiking statistics and waveform features ^35^.

### Spiking data analyses and statistics

All behavioral and spiking data were processed in MATLAB and exported to R for statistical analyses. To determine if CeA neurons were responsive to rewards we extracted the firing rate (number of spikes divided by window size) of within the reward window (0:2s after arrival at the feeder) against an approach window (−3:-1s) on each trial to create two distributions. Similar to Fraser et al. (2023), a 2-tailed Wilcoxon test (alpha = 0.05) was used to determine if a neuron was modulated by the reward on a given trial type and pooled for an overall response to rewards or their features (flavor, magnitude, or, during Reacquisition sessions, or outcome devaluation assignment) ^13^. For each neuron a ‘reward response score’ was calculated for each trial type as the firing rate in the reward window divided by the firing rate in the approach window. For visualizing the response patterns of CeA neurons to rewards, peri-event time histograms were generated using 50ms bins and were smoothed with a Gaussian (200ms). All figures were generated in MATLAB or R and stylized in Adobe Illustrator.

## Acknowledgements

KAA and KSS designed the study, KAA, EST, and JAP collected the data, JEC and JJS provided technical assistance, KAA and JEC analyzed the data, and KAA and KSS wrote the paper.

## Funding

This work was supported by NSF IOS1557987 (KSS) and NIH F99NS115270 (KAA).

## Disclosures

The authors have no competing interests to declare.

## Notes

### Competing Interest Statement

The authors have declared no competing interest.

## References

1. Balleine, B. W. & Dickinson, A. Goal-directed instrumental action: contingency and incentive learning and their cortical substrates. Neuropharmacology 37, 407–419 (1998).

2. Amaya, K. A. & Smith, K. S. Neurobiology of habit formation. Current Opinion in Behavioral Sciences vol. 20 145–152 Preprint at 10.1016/j.cobeha.2018.01.003 (2018).

3. Balleine, B. W. The Meaning of Behavior: Discriminating REFlex and Volition in the Brain. Neuron 104, 47–62 (2019).

4. Malvaez, M. Neural substrates of habit. J. Neurosci. Res. 98, 986–997 (2020).

5. Malvaez, M. & Wassum, K. M. Regulation of habit formation in the dorsal striatum. Curr Opin Behav Sci 20, 67–74 (2018).

6. Vandaele, Y. & Ahmed, S. H. Habit, choice, and addiction. Neuropsychopharmacology 46, 689–698 (2021).

7. Gasbarri, A., Pompili, A., Packard, M. G. & Tomaz, C. Habit learning and memory in mammals: behavioral and neural characteristics. Neurobiol. Learn. Mem. 114, 198–208 (2014).

8. Lingawi, N. W. & Balleine, B. W. Amygdala central nucleus interacts with dorsolateral striatum to regulate the acquisition of habits. J. Neurosci. 32, 1073–1081 (2012).

9. Warlow, S. M., Naffziger, E. E. & Berridge, K. C. The central amygdala recruits mesocorticolimbic circuitry for pursuit of reward or pain. Nat. Commun. 11, 2716 (2020).

10. Robinson, M. J. F., Warlow, S. M. & Berridge, K. C. Optogenetic excitation of central amygdala amplifies and narrows incentive motivation to pursue one reward above another. J. Neurosci. 34, 16567–16580 (2014).

11. Tom, R. L., Ahuja, A., Maniates, H., Freeland, C. M. & Robinson, M. J. F. Optogenetic activation of the central amygdala generates addiction-like pREFerence for reward. Eur. J. Neurosci. 50, 2086–2100 (2019).

12. Giovanniello, J. R. et al. A dual-pathway architecture enables chronic stress to promote habit formation. bioRxivorg (2023) doi:10.1101/2023.10.03.560731.

13. Fraser, K. M. et al. Encoding and context-dependent control of reward consumption within the central nucleus of the amygdala. bioRxivorg (2023) doi:10.1101/2023.06.28.546936.

14. Smith, K. S. & Graybiel, A. M. A dual operator view of habitual behavior REFlecting cortical and striatal dynamics. Neuron 79, 608 (2013).

15. Jin, X. & Costa, R. M. Start/stop signals emerge in nigrostriatal circuits during sequence learning. Nature 466, 457–462 (2010).

16. Graybiel, A. M. The basal ganglia and chunking of action repertoires. Neurobiol. Learn. Mem. 70, 119–136 (1998).

17. Jog, M. S., Kubota, Y., Connolly, C. I., Hillegaart, V. & Graybiel, A. M. Building neural representations of habits. Science 286, 1745–1749 (1999).

18. Barnes, T. D., Kubota, Y., Hu, D., Jin, D. Z. & Graybiel, A. M. Activity of striatal neurons REFlects dynamic encoding and recoding of procedural memories. Nature 437, 1158–1161 (2005).

19. Smith, K. S. & Graybiel, A. M. Habit formation coincides with shifts in reinforcement representations in the sensorimotor striatum. J. Neurophysiol. 115, 1487–1498 (2016).

20. Seiler, J. L. et al. Dopamine signaling in the dorsomedial striatum promotes compulsive behavior. Curr. Biol. 32, 1175-1188.e5 (2022).

21. van Elzelingen, W. et al. Striatal dopamine signals are region specific and temporally stable across action-sequence habit formation. Curr. Biol. 32, 1163–1174.e6 (2022).

22. Gremel, C. M. & Costa, R. M. Orbitofrontal and striatal circuits dynamically encode the shift between goal-directed and habitual actions. Nat. Commun. 4, 2264 (2013).

23. Thorn, C. A., Atallah, H., Howe, M. & Graybiel, A. M. Differential dynamics of activity changes in dorsolateral and dorsomedial striatal loops during learning. Neuron 66, 781–795 (2010).

24. Steinberg, E. E. et al. Amygdala-Midbrain Connections Modulate Appetitive and Aversive Learning. Neuron (2020) doi:10.1016/j.neuron.2020.03.016.

25. Esber, G. R., Torres-Tristani, K. & Holland, P. C. Amygdalo-striatal interaction in the enhancement of stimulus salience in associative learning. Behav. Neurosci. 129, 87–95 (2015).

26. Corbit, L. H. & Balleine, B. W. Double dissociation of basolateral and central amygdala lesions on the general and outcome-specific forms of pavlovian-instrumental transfer. J. Neurosci. 25, 962–970 (2005).

27. Schmitzer-Torbert, N. & Redish, A. D. Neuronal activity in the rodent dorsal striatum in sequential navigation: separation of spatial and reward responses on the multiple T task. J. Neurophysiol. 91, 2259–2272 (2004).

28. Dezfouli, A. & Balleine, B. W. Habits, action sequences and reinforcement learning. Eur. J. Neurosci. 35, 1036–1051 (2012).

29. Warlow, S. M., Robinson, M. J. F. & Berridge, K. C. Optogenetic central amygdala stimulation intensifies and narrows motivation for cocaine. J. Neurosci. 37, 8330–8348 (2017).

30. Hatfield, T., Han, J. S., Conley, M., Gallagher, M. & Holland, P. Neurotoxic lesions of basolateral, but not central, amygdala interfere with Pavlovian second-order conditioning and reinforcer devaluation effects. J. Neurosci. 16, 5256–5265 (1996).

31. Faure, A., Haberland, U., Condé, F. & El Massioui, N. Lesion to the nigrostriatal dopamine system disrupts stimulus-response habit formation. J. Neurosci. 25, 2771–2780 (2005).

32. Gonzales, C. & Chesselet, M. F. Amygdalonigral pathway: an anterograde study in the rat with Phaseolus vulgaris leucoagglutinin (PHA-L). J. Comp. Neurol. 297, 182–200 (1990).

33. Vandecasteele, M. et al. Large-scale recording of neurons by movable silicon probes in behaving rodents. J. Vis. Exp. (2012) doi:10.3791/3568-v.

34. Amaya, K. A., Stott, J. J. & Smith, K. S. Sign-tracking behavior is sensitive to outcome devaluation in a devaluation context-dependent manner: implications for analyzing habitual behavior. Learn. Mem. 27, 136–149 (2020).

35. Viskontas, I. V., Ekstrom, A. D., Wilson, C. L. & Fried, I. Characterizing interneuron and pyramidal cells in the human medial temporal lobe in vivo using extracellular recordings. Hippocampus 17, 49–57 (2007).

